# HAMLET, a human milk protein-lipid complex, modulates amoxicillin-induced changes in an *ex vivo* biofilm model of the oral microbiome

**DOI:** 10.1101/2023.11.21.568068

**Authors:** NK Brar, A Dhariwal, S Shekhar, R Junges, AP Hakansson, FC Petersen

## Abstract

Challenges from infections caused by biofilms and antimicrobial resistance highlight the need for novel antimicrobials that work synergistically with antibiotics and minimize resistance risk. In this study we investigated the potential synergistic effect of HAMLET (human alpha-lactalbumin made lethal to tumor cells), a human milk protein-lipid complex and amoxicillin on microbial ecology using an *ex-vivo* oral biofilm model. HAMLET was chosen due to its multi-targeted antimicrobial mechanism, together with its synergistic effect with antibiotics on single species pathogens, and low risk of resistance development. The combination of HAMLET and amoxicillin significantly reduced biofilm viability, while each of them alone had little or no impact. Using a whole metagenomics approach, we found that the combination group promoted a most remarkable shift in overall microbial composition compared to the untreated samples. Up to 90% of the bacterial species in the combined treatment were *Lactobacillus crispatus*, a species with probiotic effects, whereas it was detected in minor fraction in untreated samples. Resistome analysis indicated no major shifts on alpha-diversity, while beta-diversity revealed distinct clustering patterns for each treatment group, signifying that each treatment group harbors a unique resistome. TEM beta-lactamase genes were detected in low proportions in all treated samples but absent in untreated samples. Our study highlights the potential of HAMLET to synergize with amoxicillin in an *ex-vivo* model of the oral microbiome and modulate the proportion of probiotic bacteria. The findings extend the knowledge on the synergistic effects of HAMLET and antibiotics from single-species studies to polymicrobial biofilms of human origin.

**Importance:** Polymicrobial infections are challenging to treat and prevent, requiring the use of antibiotics that exhibit reduced efficacy due to biofilm formation. HAMLET has recently emerged as an antimicrobial agent that can synergize with antibiotics while limiting microbial resistance. We investigated the effects of HAMLET, alone and combined with low concentrations of amoxicillin, on *ex vivo* oral biofilms to simulate complex microbial interactions observed in the oral cavity. The combination of HAMLET and amoxicillin effectively targeted polymicrobial biofilms and led to an increase in *Lactobacillus crispatus*. The potency of this combination appears to be due to the synergistic effect of HAMLET and amoxicillin. These findings underscore the potential of combining antimicrobials with different modes of action for the development of more effective strategies for preventing and treating polymicrobial infections.

## INTRODUCTION

The human oral microbiome is considered the second-largest microbial community, following the gut microbiota, in terms of both diversity and complexity (1). A broad range of odontogenic inflammatory infections, from periodontitis and peri-implantitis to posttraumatic osteomyelitis and facial cellulitis, have been associated with oral biofilms featuring polymicrobial communities (2–7). These biofilms are difficult to treat because of their intrinsic antibiotic tolerance and resistance to the host’s immune system. Additionally, the oral cavity harbors the highest abundance of antimicrobial resistance genes (ARGs) in the entire human body, surpassing the abundance of ARGs in the gut (8). As such, the oral cavity serves as a significant potential source for the dissemination of antibiotic resistance (9, 10). Combining different therapies with potential synergistic antimicrobial activities has emerged as a novel strategy to overcome the challenges posed by polymicrobial infections (11). One particular combination that has demonstrated promising results is the utilization of HAMLET (Human Alpha-lactalbumin Made Lethal to Tumor Cells), a protein-lipid complex, in conjunction with antibiotics.

HAMLET is a complex comprised of alpha-lactalbumin and oleic acid that has demonstrated potent cancer cell killing capabilities, while sparing healthy, differentiated cells, rendering it a promising potential therapeutic. (12–14). Additionally, HAMLET exhibits antimicrobial properties against key human pathogens. Although mostly active against gram-positive bacteria, such as *Streptococcus pneumoniae* and *Streptococcus pyogenes*, HAMLET has also shown bactericidal effects on selected gram-negative species, such as *Haemophilus influenzae* and *Moraxella catarrhalis* (15). Notably, HAMLET’s bactericidal activity has not been detected in other gram-negative pathogens, including *Escherichia coli*, *Klebsiella pneumoniae*, *Pseudomonas aeruginosa*, *Haemophilus parainfluenzae*, and *Enterobacter cloacae* (15–21). When used in combination with antibiotics, HAMLET has demonstrated the ability to lower the minimum inhibitory concentration (MIC) of methicillin for methicillin resistant *Staphylococcus aureus* (MRSA) strains, bringing them within the sensitive range. HAMLET augment also the efficacy of selected antibiotics against antibiotic-resistant bacterial strains such as *S. pneumoniae* and *M. tuberculosis* (15, 18, 19).

Among the most commonly prescribed antibiotics in primary healthcare settings and for odontogenic infections is amoxicillin (22). This broad-spectrum β-lactam antibiotic is a modified form of penicillin with an extra amino group. Its mechanism of action involves disrupting peptidoglycan cross-linking in the bacterial cell-wall. Amoxicillin inactivates and kills pathogens by binding to penicillin-binding-proteins (PBPs) located on the bacterial membrane (22–24). However, its efficacy against polymicrobial biofilms such as in the oral cavity can be limited due to a number of factors, including the formation of a protective barrier that can prevent antibiotics from effectively reaching and killing the bacteria within the biofilms. Further, the production of beta-lactamases by members of microbial communities can reduce the concentration of active amoxicillin available. In combination with HAMLET, other beta-lactam antibiotics has shown synergistic effects against both *S. pneumoniae* and MRSA biofilms (15, 18). This suggests that the inclusion of HAMLET in combination with amoxicillin may have potential as an effective strategy for treating polymicrobial biofilms.

In this study, we used an *ex vivo* oral microbiome model to provide a relevant testbed for investigating the effects of HAMLET and amoxicillin on microbial ecology. Our findings indicate that the combination of amoxicillin and HAMLET act synergistically to inhibit bacterial viability in polymicrobial biofilms. Furthermore, the combination at low concentrations influenced the microbial ecology of the oral microbiome, leading to a proportional increase in bacterial species exhibiting probiotic characteristics.

## METHODS

### Sample collection

The research followed the ethical principles directed in the Declaration of Helsinki and received approval from the National Regional Ethics Committee (REK20152491) for studies involving human samples. Eight participants were instructed to brush their teeth following breakfast and to abstain from food or drink for a minimum of two hours before providing saliva samples. Additionally, they rinsed their mouths three times with water, 10 minutes before saliva collection. Non-stimulated saliva was collected from eight participants, and these samples were centrifuged at 6000 x g for 5 minutes at 4°C. This centrifugation step effectively precipitated larger debris and eukaryotic cells. The resulting supernatant was pooled and utilized as the inoculum in the human oral microbiome biofilm model, as described below.

A second centrifugation was conducted to obtain cell-free saliva by spinning down the samples at 10 000 x g for 7 minutes at 4°C. The upper fraction was used to coat the bottom of the wells prior to biofilm growth in a process termed as ‘pellicle formation’ to mimic the establishment of an oral biofilm (25).

### HAMLET production

HAMLET was produced in three steps; : 1) purification of alpha-lactalbumin from human milk, 2) converting native alpha-lactalbumin to partially unfolded protein in the presence of oleic acid (C18:1) and 3) dialysis and lyophilization as previously described (14, 21).

### The human oral microbiome biofilm model

We utilized a previously established *ex vivo* biofilm model designed to preserve a highly reproducible diversity of species and metabolic activity within the human oral microbiome (25, 26). In summary, SHI media was pre-reduced for four hours under anaerobic conditions, characterized by a carbon dioxide level of 5%, balanced with nitrogen. SHI media was prepared as previously described (27). The pooled saliva samples were added at a ratio of 2 µl of saliva per mL SHI medium. These were allotted into the wells of a 24-well plate, with each well containing 1 mL of the mixture. The plate was then incubated within an anaerobic chamber at 37°C for 24 hours.

After this incubation period, the supernatant was removed and replaced with fresh SHI medium to support the pre-formed oral biofilms. In the first set of experiments, the samples were either left untreated (control), or treated with amoxicillin ranging from 0-200 µg mL^-1^ (Sigma-Aldrich). In the second set of experiments, the preformed biofilms were not treated (control), treated with amoxicillin 0.1 µg mL^-^1, HAMLET ranging between 125-250 µg mL^-1^, or with a combination treatment composed of HAMLET ranging between 125-250µg mL^-1^in conjunction with amoxicillin at 0.1 µg mL^-^1. The stock solution of amoxicillin (2 mg/mL in distilled water) and HAMLET (5 mg/mL in phosphate-buffered saline (PBS)) were appropriately diluted in SHI medium before adding to the biofilms.

Following an incubation period of another 24 hours, the oral biofilms were washed with PBS, followed by suspension in 1 mL of PBS. Glycerol (20%) was added to the samples before they were archived and stored at −80°C.

### Oral biofilm viability assay

To evaluate the viability of the biofilms, samples obtained from both the control and the treatment groups, were subjected to a ten-fold dilution series. Subsequently, 20 µL of each dilution was plated onto SHI agar plates. These plates were then incubated for 48 hours at 37°C within an anaerobic chamber. Then the number of colony forming units per milliliter (CFUs mL^-1)^ was calculated, and represented as log 10-transformed values.

### DNA extraction

Bacterial DNA was extracted using the MasterPure^TM^ Gram Positive DNA Purification Kit (Epicentre, Madison, WI, USA), following the manufacturer’s established protocol. Subsequently, the precipitated DNA was resuspended in 35 µl milliQ water. To assess the quality and quantity of the extracted DNA, NanoDropTM 2000c spectrophotometer (Thermo Fisher Scientific, Waltham, MA, USA) was used for initial evaluation. This was followed by quantification using Qubit TM 4 Fluorometer (Thermo Fisher Scientific, Waltham, MA, USA) to yield precise measurements of the DNA’s concentrations.

### DNA library preparation and sequencing

The preparation of the DNA libraries was executed with the Illumina DNA Prep (M) kit, <u>(Illumina, Inc., San Diego, CA, USA</u>), in strict adherence to the manufacturer’s protocol. To assess the quality and concentration of the DNA library, initial measurements were conducted using the NanoDrop^TM^ 2000c spectrophotometer and Qubit ^TM^ 4 Fluorometer. Finally, analysis involved the utilization of a Bioanalyzer (Agilent Technologies, Santa Clara, CA, USA) using a High Sensitivity DNA kit (Agilent Technologies, Santa Clara, CA, USA).

The DNA library was obtained by resuspending it in the provided buffer. Each sample was adjusted to 500 ng DNA in a 30 µL volume using nuclease-free water.

For the metagenomic shotgun sequencing approach, services at the Norwegian Sequencing Centre (Oslo, Norway) were utilized, using the Illumina NovaSeq 6000 SP platform (Illumina, Inc., San Diego, CA, USA). The paired-end sequencing reads were generated with a corresponding read length of 150 base pairs.

### Assessment of sequencing read quality

The evaluation of sequencing read quality, both in raw and preprocessing state, was conducted utilizing FastQC tool (v.0.11.9) (28). The identification and removal of low-quality reads, as well as the elimination of adapter sequences, was achieved using Trimmomatic (v.0.39). The following parameters were used during this process: ILLUMINACLIP: Nextera PE:2:30:10 LEADING:3 TRAILING:3 SLIDING WINDOW:4:15 MINLEN:36.. The remaining high-quality reads were subjected to microbiome and resistome profiling.

### Taxonomic and resistome profiling

MetaPhlAn3 software (v.3.7.0) (29) was used to profile the bacterial composition in the oral biofilm samples and to determine their abundance at species-level using default settings. The ‘*merge metaphlan tables.py*’ script was used to merge the profiled metagenomes into an abundance table. To detect the hits to known Antibiotic Resistance Genes (ARGs), “high quality” paired-end reads were mapped against the Comprehensive Antibiotic Resistance Database (CARD) (v.3.2.2) (30, 31) by using the KMA alignment tool (1.4.12) (32) with parameters: *-ipe, -tmp*, *-1t1*, *-and*, *-apm f*, *-ef*. The list of detected ARGs was filtered to include only those with a minimum threshold of 80% identity between the query and reference gene over at least 80% of the reference gene length.

### Downstream analysis

Two key software tools were used to conduct comprehensive exploration, analysis, and visualization of the microbiome and resistome count data: MicrobiomeAnalyst (33, 34) and ResistoXplorer (34).

For graphical representation and statistical analysis, GraphPad Prism (Prism 9 and 10 software) as well as the R programming (version 4.2.1) were utilized. Alpha-diversity was calculated using the Shannon and Chao1 diversity indexes at species level, as well as ARG level. The top 10 most abundant features of the microbiome (species) and resistome (ARGs) data were plotted using *aggregate top taxa* and plotting functions of the microbiome R package (35–37). For Beta-diversity, Aitchison distance metric on centered log-ratio (CLR) transformed counts were performed using the *transform* and *ordinate* (RDA) function of the microbiome and phyloseq R packages. The resulting data was visualized as compositional principal component analysis (PCA) ordination plot using the *plot_ordination* function of the phyloseq package.

Pairwise comparisons of log-fold changes in the abundance of microbial species and ARGs between different groups were performed using DESeq2 (38). In order to account for multiple testing, Benjamini-Hochberg (BH) procedure was employed to adjust the results (adjusted p-values).

In case where “one-way analysis of variance” (ANOVA) was conducted, the results were adjusted for multiple comparisons using the Dunnett’s multiple comparison test. Adjusted p-values lower than 0.05 were considered statistically significant.

## RESULTS

### Dose-dependent effects of amoxicillin on oral biofilms

Pre-formed oral biofilms in new fresh SHI media were initially subjected to varying concentrations of amoxicillin, ranging from 0-200 µg/ml (**Figure 1**). The inoculum was prepared using pooled saliva obtained from eight donors. Notably, when exposed to low amoxicillin concentration within the range of 0.025-0.1 µg/ml, a slight increase in biofilm viability was observed in comparison to the negative control. The peak of this increase was remarkably evident under treatment with 0.1 µg/ml amoxicillin. However, as the amoxicillin concentration exceeded 0.1 µg/ml, a contrasting effect was observed where biofilm viability was gradually inhibited. The reduction in viability continued until the highest amoxicillin concentration was reached, at which point viable cells were almost undetectable.

**Figure 1:**
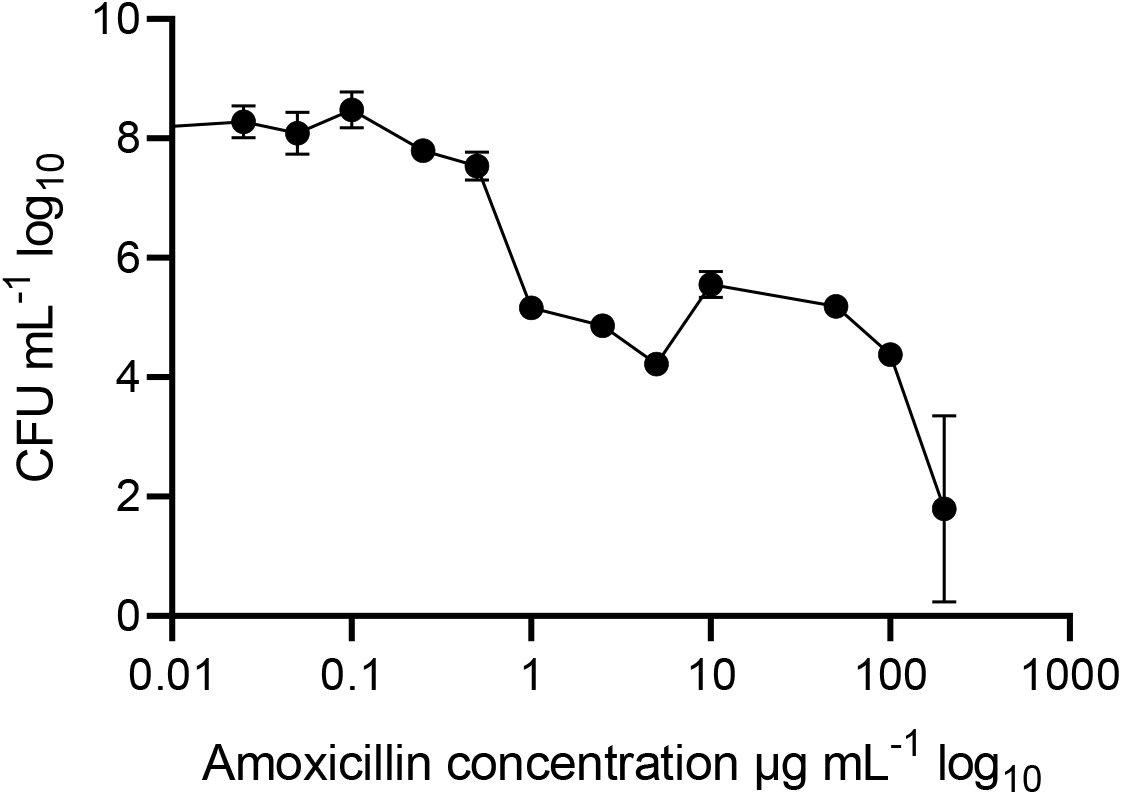
Oral biofilm treated with different amoxicillin concentrations. Number of viable cells in the polymicrobial biofilm community, as determined by colony-forming units, counted on SHI agar plates. The data are shown for triplicate experiments as mean± SE.

### The Efficacy of HAMLET and low concentrations of amoxicillin on oral biofilms

To evaluate the impact of HAMLET, both as a standalone treatment and in combination with amoxicillin at low concentrations, pre-formed oral biofilms were subjected to two different HAMLET concentrations, alone or in combination with 0.1 µg/ml of amoxicillin for 24 hours. Bactericidal activity with HAMLET alone was observed with a concentration of 250 µg/ml (Figure 2A) In context, neither HAMLET at 125 µg/ml alone nor amoxicillin alone, when assessed in comparison to the negative control, displayed any significant reduction in bacterial cell viability. However, oral biofilm viability showed to be affected by the combination treatment of HAMLET at 125 µg/ml and 0.1 µg/ml amoxicillin, leading to significant decrease in bacterial viability compared to untreated samples (Figure 2B).

**Figure 2:**
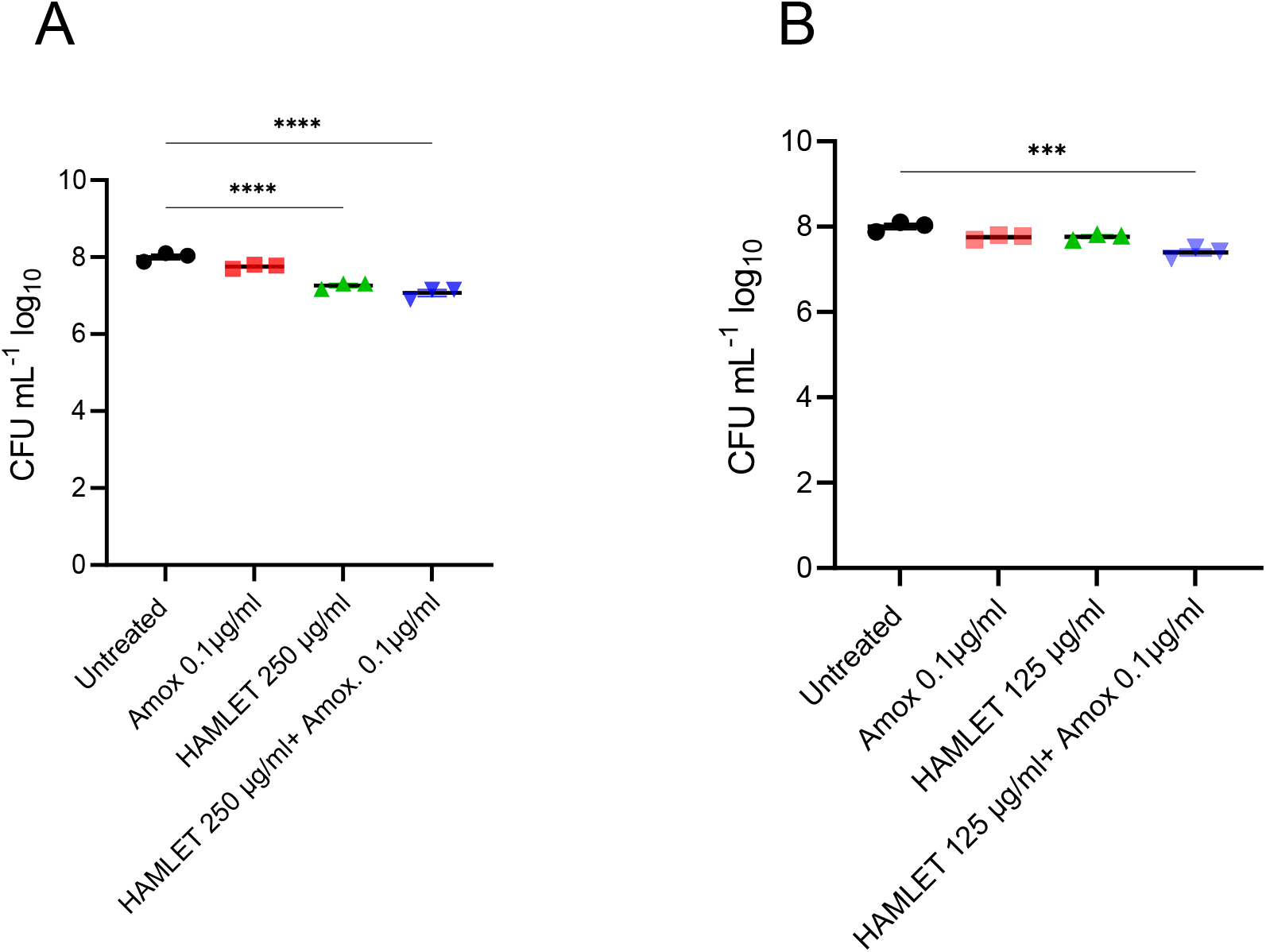
The effect of HAMLET alone or in combination with low amoxicillin concentration on oral biofilm community. (A) HAMLET concentration 250 µg/mL and (B) HAMLET concentration 125 µg/mL. Numbers of viable cells in the community are determined by colony-forming units. All results are based on three independent experiments with triplicate samples. Data are shown as mean± SE. *** P<0.001. **** P< 0.0001. One-way ANOVA followed by Dunnett’s multiple comparison test.

### Impact of HAMLET alone or in combination with amoxicillin: Oral microbiome ecology

A total of eight samples, representing two samples from each treatment group, underwent shotgun metagenomic sequencing. This analysis resulted in the generation of approximately 90.2 million paired reads per sample after quality filtering, yielding an average of 11.3 million reads per sample (with a minimum of 6.8 million and a maximum of 18.5 million reads per sample).

The metagenomic analysis provided insights into the effect of HAMLET, administrated either as a standalone treatment or in combination with amoxicillin, on the ecology of the oral microbiome. To assess changes in alpha diversity, metrics such as Chao1 index, which quantifies only microbial richness and Shannon index, which accounts for both richness and evenness (abundance) were employed. At species level, no significant changes in either alpha diversity index were observed in treatment samples when compared to the untreated control (Figure 3A**, B)**. Beta diversity analysis revealed increased variability in the biological replicates subjected to antibiotic treatment, contrasting with the biological replicates from untreated or HAMLET solely treated samples (Figure 3C).

**Figure 3:**
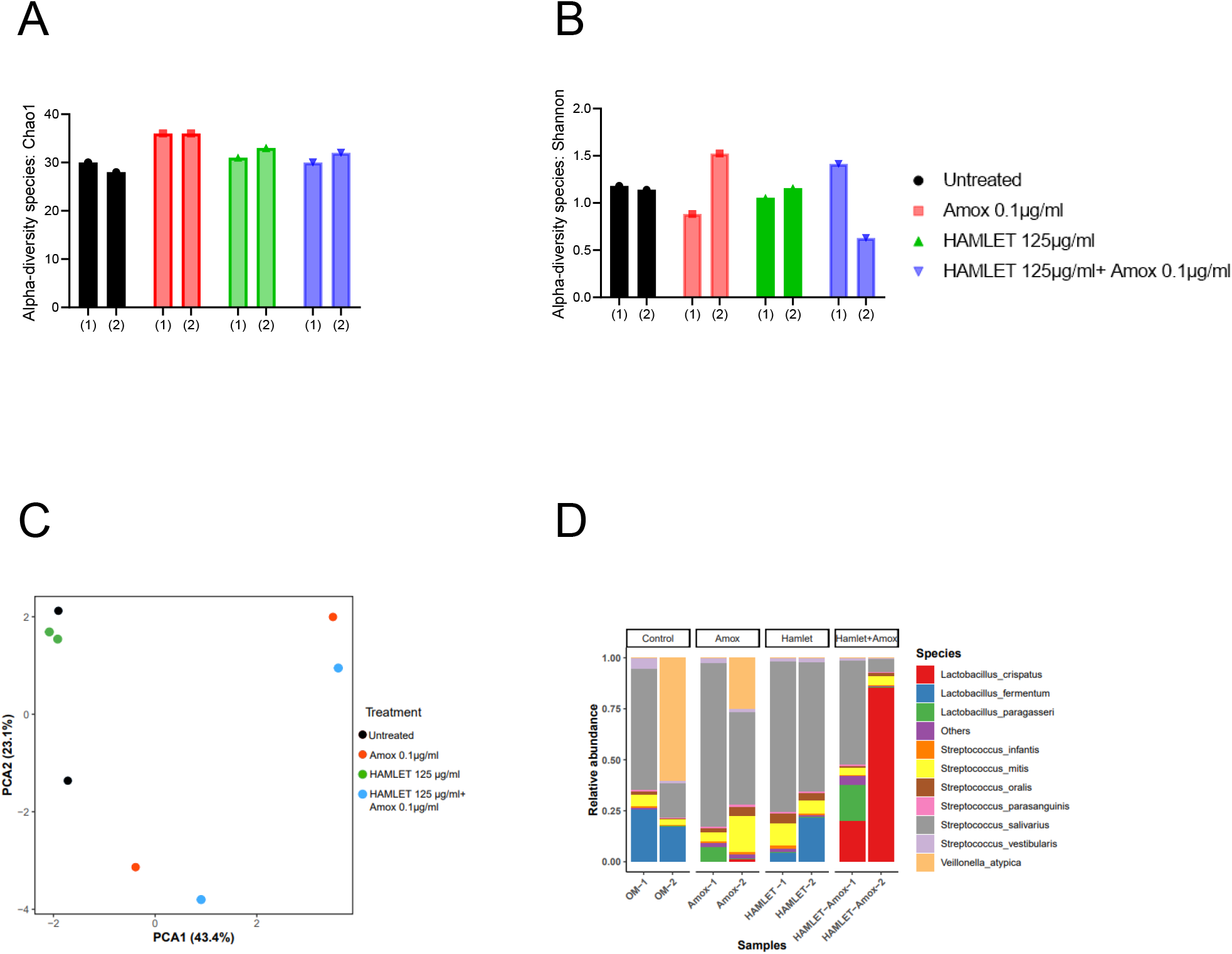
Ecological impact of HAMLET alone or in combination with amoxicillin on oral biofilm community. (A and B) Alpha-diversity on species level measured by (A) Chao1 index indicate the total richness and (B) Shannon index which indicate richness and evenness. (C) A principal component analysis plot (PCA) with Aitchison distance illustrating beta-diversity. (D) Stacked bar plots illustrate the relative abundance of all replicates for 10 most abundant species. All results are based on two biological replicates from the same day.

In total, 44 bacterial species spanning eight bacterial genera across all the samples were identified **(Supplementary Table 1)**. Despite biological sample variation, alterations in the relative abundance of taxonomic composition were evident in the treatment groups compared to the negative control **(Supplementary Table 2)**.

Analyzing the taxonomic composition at the species level revealed the emergence of new species in the amoxicillin, HAMLET and HAMLET combined with amoxicillin treated biofilm groups (Figure 3D). In comparison to the untreated samples, *Streptococcus salivarius* emerged as the dominant species in both biological replicates, while *Lactobacillus fermentum* decreased significantly in amoxicillin treated samples (**Figure S1**). Both *L. fermentum* and *S. salivarius* reduced in proportion when subjected to the combination of HAMLET and amoxicillin. In contrast, *Lactobacillus crispatus* increased significantly in proportion and dominated in the combination treatment group (**Figure S1**). *Lactobacillus paragasseri* was detected in at least one replicate of samples treated with amoxicillin, either alone or in combination with HAMLET. Predominant colonizers were gram-positives, with the exception of two samples that had more than 20% *<u>Veillonella atypica</u>*. However, these were in replicates that belonged to different treatment groups.

Furthermore, the presence of the pathogen *Streptococcus pneumoniae* was noted in all samples, with an increase in amoxicillin-treated samples compared to the untreated control **(Supplementary Table 1)**.

### Impact of HAMLET alone or in combination with amoxicillin on Oral resistome

Across all the eight samples, a total of 123,350 paired reads were annotated as antimicrobial resistant genes (ARGs). On average, there were 15,418 reads per sample, with a minimum count of 3,625 and a maximum count of 24,976. In total, 22 distinct ARGs associated with seven antibiotic drug classes and four antibiotic resistance mechanisms: antibiotic efflux, antibiotic inactivation, antibiotic target alteration, and antibiotic target protection were identified **(Figure S2, Supplementary Table 3).**

Alpha-diversity, as measured by Chao 1 and Shannon indexes exhibited no major changes in the treatment groups when compared to the untreated control (Figure 4A, B). However, beta-diversity analysis revealed distinct clustering patterns of biological samples within each treatment group, signifying that each treatment group harbors a unique resistome (Figure 4C).

**Figure 4:**
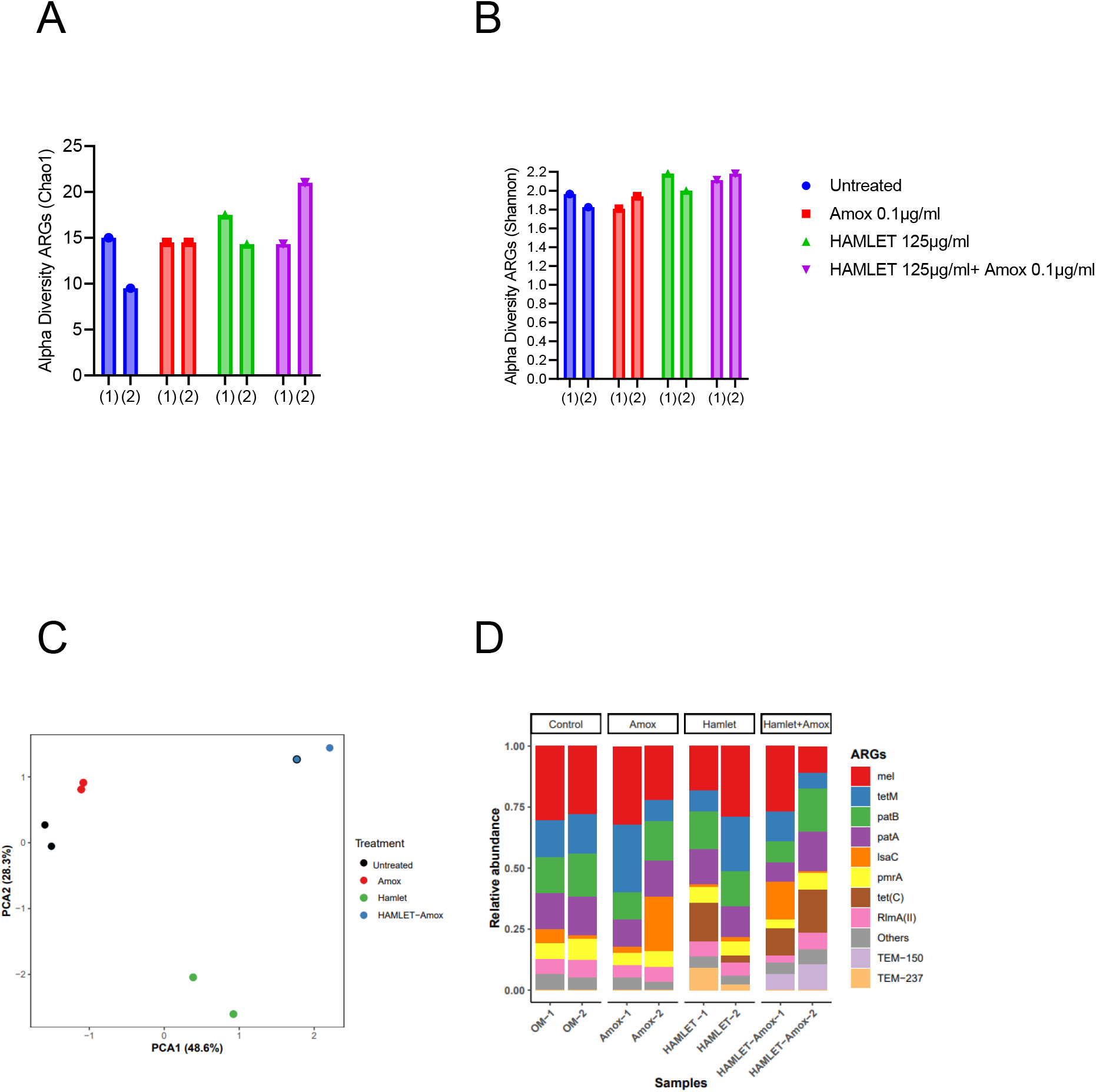
Impact of HAMLET alone or in combination with amoxicillin on oral resistome. (A and B) Alpha-diversity on ARG level measured by (A) Chao1 index indicates the total richness and (B) Shannon index which indicates richness and evenness. (C) Principal component analysis (PCA) ordination plot with Aitchison distance illustrating beta-diversity. (D) Stacked bar plots illustrate the relative abundance of all replicates for the 10 most abundant ARG’s. All results are based on two biological replicates from the same day.

ARGs associated with all four antibiotic resistance mechanisms were detected in all treatment groups, with the highest relative abundance observed in ARGs related to antibiotic target protection and antibiotic efflux (Figure S2). The three most prevalent classes of ARGs in all treatment groups included fluoroquinolone, tetracycline, and macrolide-lincosamide-streptogramin (MLS). Although there was a modest increase in the beta-lactam ARG class in HAMLET and the combination treatment with HAMLET and amoxicillin. (Figure S2).

Regarding specific ARGs, despite some variations between the replicates, *mel*, *patA*, *patB*, *pmrA*, *RImA(II)*, and *tetM* genes were detected in high abundance across all treatment groups (Figure 4D and Supplementary Table 3). Furthermore, the relative abundance of the *tet(C)* gene showed an increase in the HAMLET and HAMLET combined with amoxicillin-treated samples. For the beta-lactam antibiotic resistance genes, TEM genes were detected in all treated samples, although they comprised a low proportion of all ARGs.

## DISCUSSION

With the treatment challenges of infections caused by biofilms and the growing global issue of antimicrobial resistance, there is an increased interest in identifying novel antimicrobials that can work synergistically with antibiotics while lowering the likelihood of microbial resistance (39, 40). Here, we investigated the combined usage of HAMLET and amoxicillin. HAMLET was specifically chosen due to its unique multi-targeted antimicrobial mechanism including inhibition of glycolytic pathways (20). Other promising properties of HAMLET such as, lack of resistance development in studies with *S. pneumoniae* and *S. aureus* (*15, 18*), and established low-or non-toxic profile in prior animal and human studies investigating its potential as an anticancer drug (41–44).

Our focus was on polymicrobial biofilm communities, an area that has received less attention compared to single bacterium in biofilms (45, 46). Our results revealed that neither HAMLET nor amoxicillin individually in our comparative analysis had a significant effect on the overall cell viability of the polymicrobial community compared with the untreated control at the chosen concentrations. However, their combination resulted in a significant reduction in biofilm viability, indicating a synergistic effect. Furthermore, through metagenomic analysis, our data suggested that this combination may skew the polymicrobial community towards populations with potential probiotic effects, thereby representing a potential new approach on managing polymicrobial biofilms.

One of the most studied probiotic bacteria are lactobacilli. Several species in this genus have been shown to have beneficial effects, including improving gut and oral health, boosting the immune system, aiding in the digestion of lactose, and reducing the risk of certain infections (47–51). *Lactobacillus fermentum* was the dominant species of lactobacilli in the non-treated control samples, comprising approximately 25% of the microbiome. These were practically absent in amoxicillin treated samples. It was therefore interesting that the combination of HAMLET and low-amoxicillin concentration in our study resulted in a microbiome dominated by lactobacilli. *Lactobacillus crispatus,* in particular, was among those that increased in abundance from very low detected levels in the non-treated control and single treatments with HAMLET or amoxicillin to up to 90% with the combination of the agents. This species is known for its probiotic and antimicrobial properties (52), and has been associated with oral health, particularly in the context of dental and periodontal diseases (53–55) These results highlight the potential of the combination of HAMLET and amoxicillin to modulate the composition of the microbiome towards a community enriched in probiotic bacteria, compared to samples treated with amoxicillin alone.

Although our study primarily aimed to investigate the combined effects of HAMLET and amoxicillin, the observations on HAMLET alone are also of relevance. We initially tested two concentrations of HAMLET for its potential effect on cell viability. From these, we chose the lowest concentration of HAMLET for the metagenomics studies to underscore its potential synergistic effect with amoxicillin. Although at this low dosage HAMLET alone had no discernible effects on the overall number of viable bacteria, the metagenomics analysis indicated potential changes in the microbiome composition with increased relative abundance of *S. salivarius*. It is possible that the changes by HAMLET, alone or in combination with amoxicillin, are a result of its influence on glycolytic pathways. Previous studies have shown that HAMLET binds to and inactivates two key glycolytic enzymes, fructose-bisphosphate aldolase and glyceraldehyde-3-phosphate dehydrogenase (GAPDH) (20). However, the mechanism of HAMLET’s antimicrobial effect is not yet fully understood.

In the case of antibiotics, it is known that at low concentrations bacteria can sense antibiotics as a stress, and rather than eliminating them, these low concentrations may promote stress responses that can favor overall survival (10, 56–58). This adds a level of complexity that needs to be considered in interpreting results using polymicrobial communities. Small changes in one microbial species caused by low concentrations of antimicrobials can trigger major ecological shifts in the community due to interdependent and nonlinear interactions between microbial species.

In our study, we observed that *TEM* genes encoding beta-lactamases (59) were present in samples exposed to HAMLET alone, amoxicillin alone or in combination with amoxicillin. While the increase in abundance of this gene in samples exposed to amoxicillin could potentially be linked to survival mechanisms in the presence of this beta-lactam antibiotic, it is difficult to explain the presence of this beta-lactamase in the HAMLET group alone. We also observed an increase in the abundance of *(tet)C* in both the groups treated with HAMLET alone and in combination with amoxicillin, despite tetracycline not being used in the study. Such increases in antibiotic resistance genes in response to the presence of low concentrations of antimicrobials are frequently reported in metagenomic studies and are often not solely attributed to the co-carriage of different antibiotic resistance genes in mobile genetic elements (60–62). Instead, this phenomenon is reflective of the complex and intricate ecological dynamics within microbial communities, as discussed above.

In conclusion, our results highlight the potential of HAMLET as a synergistic antimicrobial agent when combined with amoxicillin. The significant shift in the oral microbiome towards an increase in *Lactobacillus crispatus*, a potential probiotic, by the combined agents presents a promising strategy for combatting polymicrobial infections and reducing the burden of antibiotic resistance. The findings of our study suggest that this approach may contribute to the development of more effective strategies for combating drug-resistant polymicrobial infections and underscore the importance of continued research in this area.

## ACKNOWLEDGMENTS

The sequencing service was provided by the Norwegian Sequencing Centre (www.sequencing.uio.no). The sequencing analysis and storage were performed on SAGA & NIRD resources provided by Sigma2-the National Infrastructure for High Performance Computing and Data Storage in Norway.

We thank the Research Council of Norway (RCN) (project numbers: 273833, 274867, and 322375), and the Olav Thon Foundation for funding support.

## AUTHOR CONTRIBUTION

Conceptualization of the project was by FCP and APH. All authors contributed to the design of experiments. Laboratory work was by NKB, APH, FCP. Downstream metagenomics analysis was by NKB and AD. First drafting of manuscript by NKB, FCP, AD. All authors contributed to critical review of data and writing the final manuscript.

## COMPETING OF INTERESTS

No potential conflict of interest was reported by the author(s).

## ADDITIONAL INFORMATION

### Ethics statement

The study was conducted in accordance with the Declaration of Helsinki and approved by the National Regional Ethical Committee (REK20152491)

## SUPPLEMENTARY FIGURES

**Figure S1:**
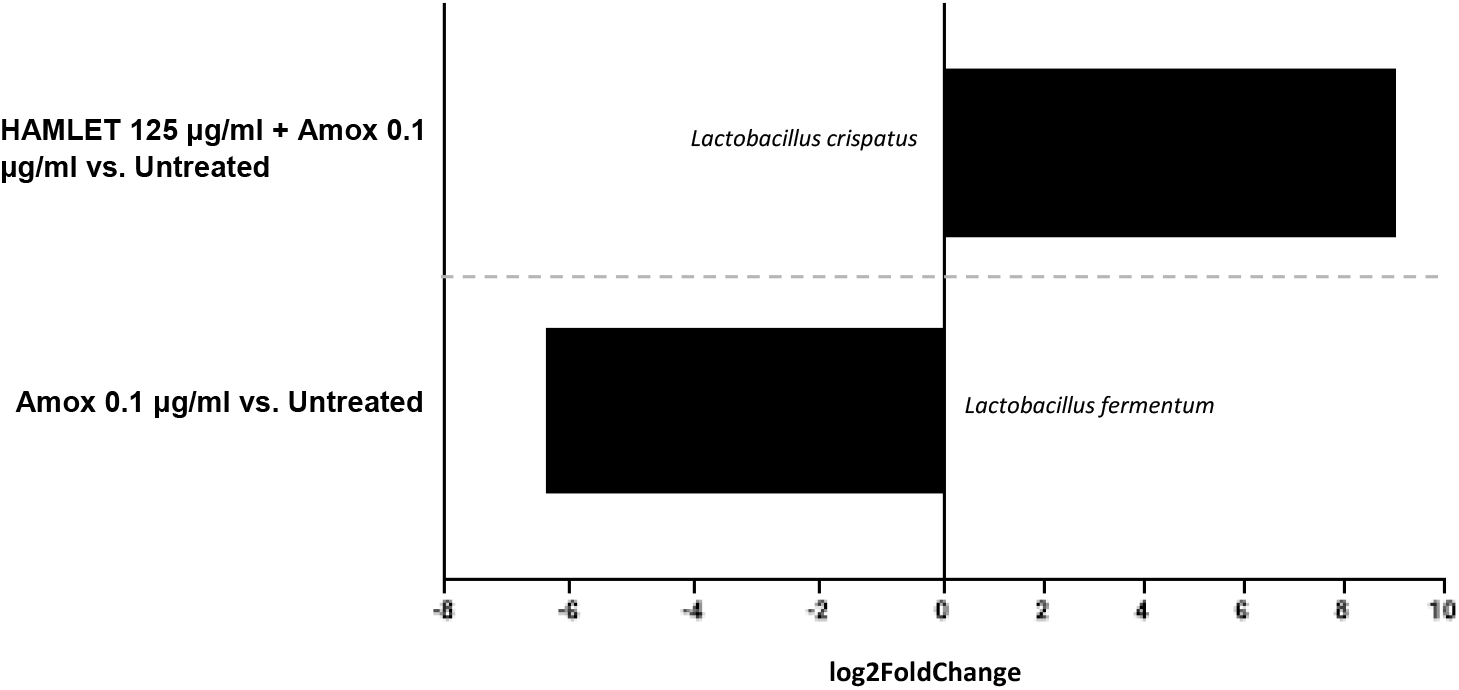
Taxa with significantly different abundance upon treatment. Bar charts illustrate the log_2_ fold change of taxa, adjusted for false discovery rate (FDR), *p*-values <0.05. (based on DESeq2).

**Figure S2:**
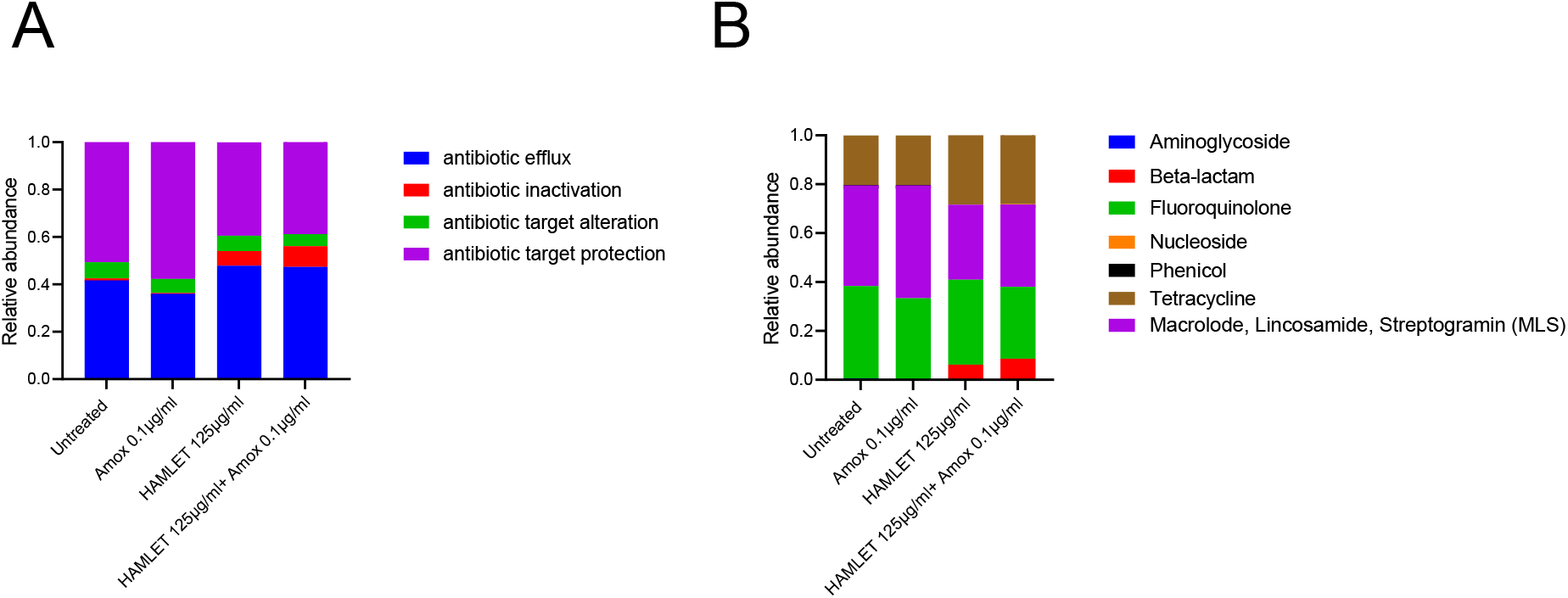
Impact of HAMLET alone or in combination with amoxicillin on oral resistome. (A and B) Stacked bar plots display the relative abundance of (A) all antibiotic mechanisms (B) all ARG classes.

## REFERENCES

1. Caselli E, Fabbri C, D’Accolti M, Soffritti I, Bassi C, Mazzacane S, et al. Defining the oral microbiome by whole-genome sequencing and resistome analysis: the complexity of the healthy picture. BMC microbiology. 2020;20(1):1–19.

2. Anju V, Busi S, Imchen M, Kumavath R, Mohan MS, Salim SA, et al. Polymicrobial infections and biofilms: clinical significance and eradication strategies. Antibiotics. 2022;11(12):1731.

3. Costa RC, Bertolini M, Costa Oliveira BE, Nagay BE, Dini C, Benso B, et al. Polymicrobial biofilms related to dental implant diseases: unravelling the critical role of extracellular biofilm matrix. Critical Reviews in Microbiology. 2023;49(3):370–90.

4. Hajishengallis G, Lamont RJ. Polymicrobial communities in periodontal disease: Their quasi-organismal nature and dialogue with the host. Periodontology 2000. 2021;86(1):210-30.

5. Dashper SG, Nastri A, Abbott PV. Odontogenic bacterial infections. Contemporary Oral Medicine: A comprehensive approach to clinical practice Cham, Switzerland: Springer Nature. 2019:819-70.

6. Gaetti-Jardim E, Landucci LF, de Oliveira KL, Costa I, Ranieri RV, Okamoto AC, et al. Microbiota associated with infections of the jaws. International journal of dentistry. 2012;2012.

7. Masters EA, Ricciardi BF, Bentley KLdM, Moriarty TF, Schwarz EM, Muthukrishnan G. Skeletal infections: microbial pathogenesis, immunity and clinical management. Nature Reviews Microbiology. 2022;20(7):385–400.

8. Carr VR, Witherden EA, Lee S, Shoaie S, Mullany P, Proctor GB, et al. Abundance and diversity of resistomes differ between healthy human oral cavities and gut. Nature communications. 2020;11(1):693.

9. Anderson AC, von Ohle C, Frese C, Boutin S, Bridson C, Schoilew K, et al. The oral microbiota is a reservoir for antimicrobial resistance: resistome and phenotypic resistance characteristics of oral biofilm in health, caries, and periodontitis. Annals of clinical microbiology and antimicrobials. 2023;22(1):37.

10. Roberts AP, Mullany P. Oral biofilms: a reservoir of transferable, bacterial, antimicrobial resistance. Expert review of anti-infective therapy. 2010;8(12):1441–50.

11. Shrestha L, Fan H-M, Tao H-R, Huang J-D. Recent strategies to combat biofilms using antimicrobial agents and therapeutic approaches. Pathogens. 2022;11(3):292.

12. HÅKANssoN A, Zhivotovsky B, Orrenius S, SABHARwAL H, Svanborg C. Apoptosis induced by a human milk protein. Proceedings of the National Academy of Sciences. 1995;92(17):8064–8.

13. Svensson M, Düringer C, Hallgren O, Mossberg A-K, Håkansson A, Linse S, et al. Hamlet—a complex from human milk that induces apoptosis in tumor cells but spares healthy cells. Integrating population outcomes, biological mechanisms and research methods in the study of human milk and lactation. 2002:125–32.

14. Svensson M, Håkansson A, Mossberg A-K, Linse S, Svanborg C. Conversion of α-lactalbumin to a protein inducing apoptosis. Proceedings of the National Academy of Sciences. 2000;97(8):4221–6.

15. Marks LR, Clementi EA, Hakansson AP. The human milk protein-lipid complex HAMLET sensitizes bacterial pathogens to traditional antimicrobial agents. 2012.

16. Alamiri F, Riesbeck K, Hakansson AP. HAMLET, a protein complex from human milk, has bactericidal activity and enhances the activity of antibiotics against pathogenic Streptococci. Antimicrobial Agents and Chemotherapy. 2019;63(12):e01193–19.

17. Hakansson AP, Roche-Hakansson H, Mossberg A-K, Svanborg C. Apoptosis-like death in bacteria induced by HAMLET, a human milk lipid-protein complex. PLoS One. 2011;6(3):e17717.

18. Marks LR, Clementi EA, Hakansson AP. Sensitization of Staphylococcus aureus to methicillin and other antibiotics in vitro and in vivo in the presence of HAMLET. PloS one. 2013;8(5):e63158.

19. Meikle V, Mossberg A-K, Mitra A, Hakansson AP, Niederweis M. A protein complex from human milk enhances the activity of antibiotics and drugs against Mycobacterium tuberculosis. Antimicrobial Agents and Chemotherapy. 2019;63(2):e01846–18.

20. Roche-Hakansson H, Vansarla G, Marks LR, Hakansson AP. The human milk protein-lipid complex HAMLET disrupts glycolysis and induces death in Streptococcus pneumoniae. Journal of Biological Chemistry. 2019;294(51):19511–22.

21. Håkansson A, Svensson M, Mossberg AK, Sabharwal H, Linse S, Lazou I, et al. A folding variant of α-lactalbumin with bactericidal activity against streptococcus pneumoniae. Molecular microbiology. 2000;35(3):589–600.

22. Akhavan BJ KN, Vijhani P. Amoxicillin 2023 [Available from: https://www.ncbi.nlm.nih.gov/books/NBK482250/.

23. PubChem Compound Summary for CID 33613, Amoxicillin: National Center for Biotechnology Information (2023); 2023 [Available from: https://pubchem.ncbi.nlm.nih.gov/compound/Amoxicillin.

24. Akhavan BJ, Khanna NR, Vijhani P. Amoxicillin. StatPearls [Internet]: StatPearls Publishing; 2021.

25. Edlund A, Yang Y, Hall AP, Guo L, Lux R, He X, et al. An in vitro biofilm model system maintaining a highly reproducible species and metabolic diversity approaching that of the human oral microbiome. Microbiome. 2013;1:1–17.

26. Edlund A, Yang Y, Yooseph S, He X, Shi W, McLean JS. Uncovering complex microbiome activities via metatranscriptomics during 24 hours of oral biofilm assembly and maturation. Microbiome. 2018;6(1):1–22.

27. Tian Y, He X, Torralba M, Yooseph S, Nelson K, Lux R, et al. Using DGGE profiling to develop a novel culture medium suitable for oral microbial communities. Molecular oral microbiology. 2010;25(5):357–67.

28. Andrews S. FastQC: a quality control tool for high throughput sequence data. Babraham Bioinformatics, Babraham Institute, Cambridge, United Kingdom; 2010.

29. Truong DT, Franzosa EA, Tickle TL, Scholz M, Weingart G, Pasolli E, et al. MetaPhlAn2 for enhanced metagenomic taxonomic profiling. Nature methods. 2015;12(10):902–3.

30. Alcock BP, Raphenya AR, Lau TT, Tsang KK, Bouchard M, Edalatmand A, et al. CARD 2020: antibiotic resistome surveillance with the comprehensive antibiotic resistance database. Nucleic acids research. 2020;48(D1):D517-D25.

31. Database TCAR. The Comprehensive Antibiotic Resistance Database CARD 2020 [Available from: https://card.mcmaster.ca/.

32. Clausen PT, Aarestrup FM, Lund O. Rapid and precise alignment of raw reads against redundant databases with KMA. BMC bioinformatics. 2018;19:1–8.

33. Chong J, Liu P, Zhou G, Xia J. Using MicrobiomeAnalyst for comprehensive statistical, functional, and meta-analysis of microbiome data. Nature protocols. 2020;15(3):799–821.

34. Dhariwal A, Chong J, Habib S, King IL, Agellon LB, Xia J. MicrobiomeAnalyst: a web-based tool for comprehensive statistical, visual and meta-analysis of microbiome data. Nucleic acids research. 2017;45(W1):W180–W8.

35. McMurdie PJ, Holmes S. phyloseq: an R package for reproducible interactive analysis and graphics of microbiome census data. PloS one. 2013;8(4):e61217.

36. Lahti L, Shetty S. microbiome R package. 2017.

37. Wickham H, Wickham H. Getting Started with ggplot2. ggplot2: Elegant graphics for data analysis. 2016:11-31.

38. Love MI, Huber W, Anders S. Moderated estimation of fold change and dispersion for RNA-seq data with DESeq2. Genome biology. 2014;15(12):1–21.

39. Basavegowda N, Baek K-H. Combination strategies of different antimicrobials: An efficient and alternative tool for Pathogen inactivation. Biomedicines. 2022;10(9):2219.

40. Kaneti G, Meir O, Mor A. Controlling bacterial infections by inhibiting proton-dependent processes. Biochimica et Biophysica Acta (BBA)-Biomembranes. 2016;1858(5):995–1003.

41. Ho JC, Nadeem A, Svanborg C. HAMLET–A protein-lipid complex with broad tumoricidal activity. Biochemical and biophysical research communications. 2017;482(3):454–8.

42. Mok KH, Pettersson J, Orrenius S, Svanborg C. HAMLET, protein folding, and tumor cell death. Biochemical and biophysical research communications. 2007;354(1):1–7.

43. Puthia M, Storm P, Nadeem A, Hsiung S, Svanborg C. Prevention and treatment of colon cancer by peroral administration of HAMLET (human α-lactalbumin made lethal to tumour cells). Gut. 2014;63(1):131–42.

44. Gustafsson L, Leijonhufvud I, Aronsson A, Mossberg A-K, Svanborg C. Treatment of skin papillomas with topical α-lactalbumin–oleic acid. New England Journal of Medicine. 2004;350(26):2663–72.

45. Joshi RV, Gunawan C, Mann R. We are one: multispecies metabolism of a biofilm consortium and their treatment strategies. Frontiers in Microbiology. 2021;12:635432.

46. Gabrilska RA, Rumbaugh KP. Biofilm models of polymicrobial infection. Future microbiology. 2015;10(12):1997–2015.

47. Chugh P, Dutt R, Sharma A, Bhagat N, Dhar MS. A critical appraisal of the effects of probiotics on oral health. Journal of functional foods. 2020;70:103985.

48. Kechagia M, Basoulis D, Konstantopoulou S, Dimitriadi D, Gyftopoulou K, Skarmoutsou N, et al. Health benefits of probiotics: a review. International Scholarly Research Notices. 2013;2013.

49. Mazziotta C, Tognon M, Martini F, Torreggiani E, Rotondo JC. Probiotics mechanism of action on immune cells and beneficial effects on human health. Cells. 2023;12(1):184.

50. Li X, Wang Q, Hu X, Liu W. Current status of probiotics as supplements in the prevention and treatment of infectious diseases. Frontiers in Cellular and Infection Microbiology. 2022;12:167.

51. Kim HS, Gilliland SE. Lactobacillus acidophilus as a dietary adjunct for milk to aid lactose digestion in humans. Journal of dairy science. 1983;66(5):959–66.

52. Terai T, Kato K, Ishikawa E, Nakao M, Ito M, Miyazaki K, et al. Safety assessment of the candidate oral probiotic Lactobacillus crispatus YIT 12319: Analysis of antibiotic resistance and virulence-associated genes. Food and Chemical Toxicology. 2020;140:111278.

53. Terai T, Okumura T, Imai S, Nakao M, Yamaji K, Ito M, et al. Screening of probiotic candidates in human oral bacteria for the prevention of dental disease. PloS one. 2015;10(6):e0128657.

54. Wang J, Liu Y, Wang W, Ma J, Zhang M, Lu X, et al. The rationale and potential for using Lactobacillus in the management of periodontitis. Journal of Microbiology. 2022;60(4):355–63.

55. Lin P-P, Hsieh Y-M, Tsai C-C. Isolation and Characterisation of Probiotics for Antagonising Cariogenic Bacterium Streptococcus mutans and Preventing Biofilm Formation. 2016.

56. Penesyan A, Paulsen IT, Gillings MR, Kjelleberg S, Manefield MJ. Secondary effects of antibiotics on microbial biofilms. Frontiers in microbiology. 2020;11:2109.

57. Preda VG, Săndulescu O. Communication is the key: biofilms, quorum sensing, formation and prevention. Discoveries. 2019;7(3).

58. Ranieri MR, Whitchurch CB, Burrows LL. Mechanisms of biofilm stimulation by subinhibitory concentrations of antimicrobials. Current opinion in microbiology. 2018;45:164–9.

59. Bush K, Bradford PA. Epidemiology of β-lactamase-producing pathogens. Clinical microbiology reviews. 2020;33(2):10.1128/cmr. 00047-19.

60. Shi Y, Zhang Y, Wu X, Zhang H, Yang M, Tian Z. Potential dissemination mechanism of the tetC gene in Aeromonas media from the aerobic biofilm reactor under oxytetracycline stresses. Journal of Environmental Sciences. 2021;105:90–9.

61. Andersson DI, Hughes D. Microbiological effects of sublethal levels of antibiotics. Nature Reviews Microbiology. 2014;12(7):465–78.

62. Partridge SR, Kwong SM, Firth N, Jensen SO. Mobile genetic elements associated with antimicrobial resistance. Clinical microbiology reviews. 2018;31(4):10.1128/cmr. 00088-17.

